# Voronota-LT: efficient, flexible and solvent-aware tessellation-based analysis of atomic interactions

**DOI:** 10.1101/2024.02.05.577169

**Authors:** Kliment Olechnovič, Sergei Grudinin

## Abstract

**Motivation:** In the fields of structural biology and bioinformatics, understanding molecular interactions is paramount. However, existing advanced geometric methods for describing interatomic contacts considering full structural context have typically demanded substantial computational resources, hindering their practical application. Given the ever-growing volume of structural data, there is an urgent need for more efficient tools for interaction analysis.

**Results:** We present Voronota-LT, a new efficient method tailored for computing Voronoi tessellation-based atom-atom contacts within the solvent-accessible surface of molecular structures. Voronota-LT constructs every interatomic contact surface directly, without pre-computing the global Voronoi diagram or Delaunay triangulation. This approach results in a method that is not only fast, but also parallelizable and capable of selectively targeting specific interface areas within molecular complexes. While offering high execution speed, Voronota-LT provides a comprehensive description of every interatomic interaction, taking full account of the relevant structural context.

**Availability and Implementation:** Voronota-LT software is freely available as both a standalone application and a C++ header-only library at https://kliment-olechnovic.github.io/voronota/expansion_lt/.

## 1. Introduction

There are multiple ways to computationally define and analyze interatomic contacts in a three-dimensional (3D) model of a molecular structure, but most studies in structural bioinformatics only rely on calculating and interpreting atom-atom distances. This is not surprising, as such analysis is usually fast and easy to implement. However, there are alternative approaches that are more descriptive. For example, the Voronoi tessellation (Voronoi, 1908) and its weighted variants that can consider atomic radii, Laguerre-Voronoi diagram (Imai et al., 1985; Aurenhammer, 1987) and additively weighted Voronoi tessellation (Goede et al., 1997; Kim et al., 2005), were known for a long time and used to investigate various properties of molecular structures (Poupon, 2004; Cazals, 2006).

While distance calculation for a pair of atoms does not depend on the other surrounding atoms, the Voronoi tessellation-based interaction analysis accounts for the structural neighbors that may affect a given contact (Esque et al., 2011). Implicit interactions with solvent can be added by constraining Voronoi cells inside the solvent-accessible surface of a molecule (Goede et al., 1997). The faces of the constrained Voronoi cells can be described quantitatively by calculating their areas after on-sphere projection (McConkey et al., 2002) or directly (Olechnovič and Venclovas, 2017). Tessellation-derived contact areas can then be used to define machine learning-based scoring functions that estimate, for example, pseudo-energy for every interaction (McConkey et al., 2003; Verdonk et al., 2016; Olechnovič and Venclovas, 2017, 2023; Igashov et al., 2021a,b). Because such contact area-based scores correspond to the partitioning of space between atoms, they can be added up to obtain to a global score value that accounts for the packing arrangement of atoms. In contrast, pairwise distance-based scoring functions produce values that are inherently non-additive from the perspective of statistical physics (Ben-Naim, 1997).

The most direct way of constraining Voronoi tessellation-derived atom-atom contacts inside the solvent-accessible surface is simply cutting Voronoi cell faces by the solvent-accessible surface. This approach is utilized for computing areas of constrained Voronoi cell faces in several algorithms that calculate volumes of intersecting balls (Edelsbrunner and Koehl, 2003; Cazals et al., 2011; Klenin et al., 2011). However, the detailed atom-atom contact areas are rarely treated as a useful product of such algorithms. Surprisingly, to the best of our knowledge, only one publicly available software tool outputs solvent-constrained tessellation-based interatomic contact areas for the analysis of molecular structures — Voronota (Olechnovič and Venclovas, 2014, 2021). There are recent testimonies to the usefulness of Voronota-computed contacts. Indeed, they were used as the basis for the best-performing method (Olechnovič et al., 2023) in the scoring challenge in the recent CASP15-CAPRI experiment (Lensink et al., 2023), and they comprised the inter-chain interface feature descriptors that were assessed to be some of the most useful for discriminating physiological from non-physiological interfaces in structures of protein complexes (Schweke et al., 2023). It was also reported that Voronota computes the global Voronoi tessellation of balls relatively very rapidly when the input is biomolecular structural data (Lee et al., 2022). Nevertheless, the total time that Voronota needs to compute contacts can still reach several seconds for a moderately-sized biomolecular structure of about 5,000 atoms.

In this work, we present a new approach to constructing Voronoi tessellation-derived atom-atom contacts for macromolecules with conservatively defined solvent-accessibility surfaces. We show how to efficiently construct tessellation-compatible contact surfaces directly, without pre-computing the Voronoi diagram or Delaunay triangulation. The new method is called “Voronota-LT” (pronounced as “Voronota lite”).

## 2. Methods

### 2.1. Definition of an atom-atom contact

Given an input molecular structure in 3D, we interpret atoms as balls of van der Waals radii. For every atom, we define a *contact sphere* around it by augmenting the atomic ball radius with the solvent probe radius (commonly set to 1.4 Å). We say that two atoms are *close enough* to have a physically relevant contact if their contact spheres intersect, that is, if a solvent probe cannot fit between the two atomic balls. When considering such two atoms outside of any neighboring structural context, we define the contact between the atoms to be the *disk* defined by the intersection of the two contact spheres (Fig. 1a). When atoms have multiple neighbors, there can be multiple contact disks that can intersect (Fig. 1b).

**Fig. 1:**
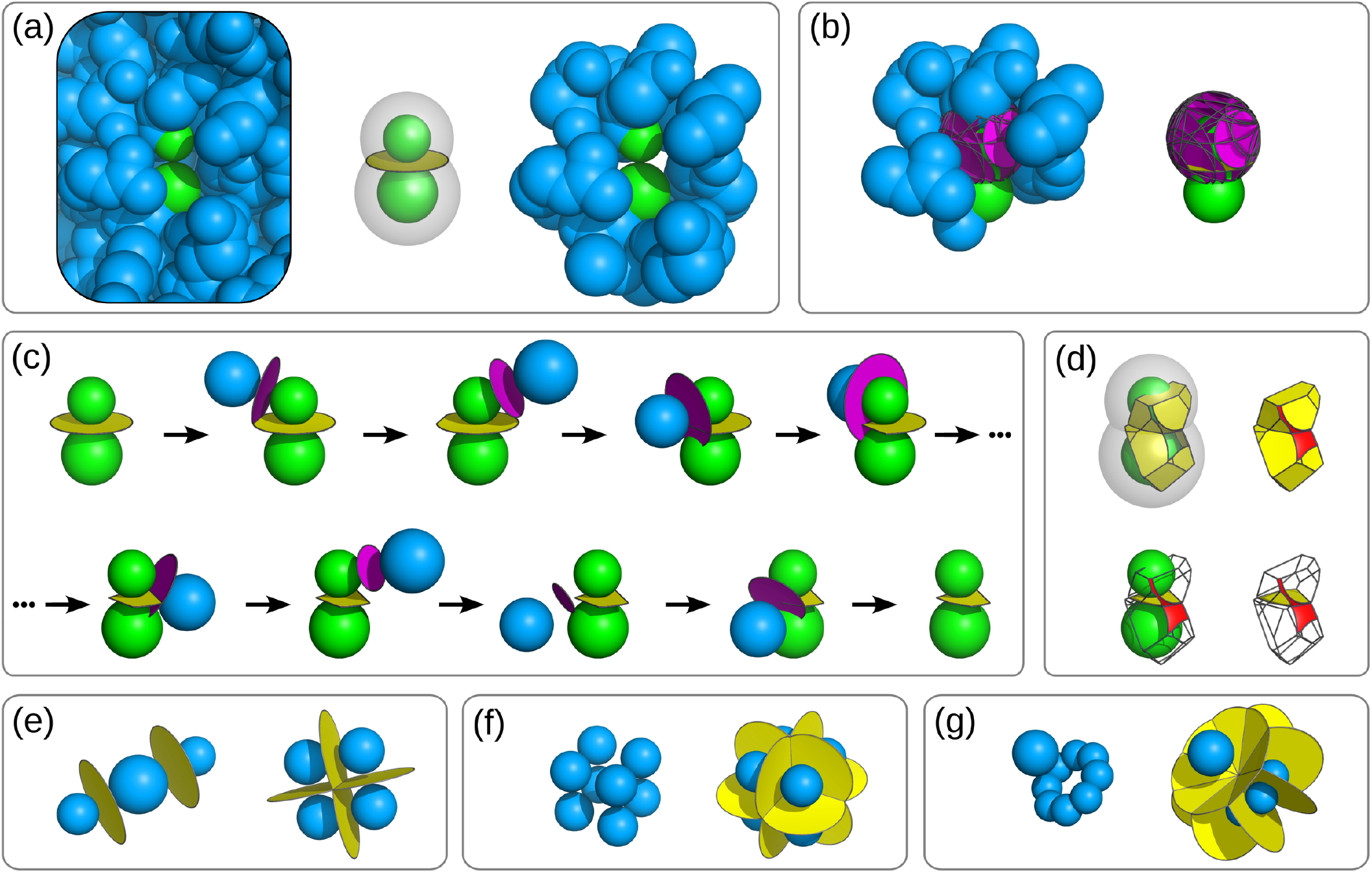
Algorithmic aspects of Voronota-LT. **(a)** A pair of possibly contacting atoms (colored green) surrounded by other atoms (colored blue) in a protein structure (left); the maximum-area contact defined as a disk produced by intersecting two probe-expanded spheres (middle); neighboring blue atoms that are close enough to have contact disks with either of the green atoms (right). **(b)** All close-enough neighbors of the first green atom (left), and the corresponding contact disks (right). **(c)** Illustration of the iterative contact cutting process, starting with the maximum-area contact defined as a disk and ending with the final contact that cannot be cut anymore. **(d)** Illustrations of the constrained Laguerre-Voronoi cells formed by all the produced contacts of the green atoms. The exposed circular boundaries of the contacts define solvent-accessible surface patches (colored red) on the probe-expanded spheres. **(e-g)** Examples of contacts constructed correctly for arrangements of balls that induce degenerate Laguerre-Voronoi cells: balls with centers on a line and on a square lattice **(e)**; balls on a cubic lattice **(f)**, balls in a rotationally symmetrical arrangement **(g)**.

Every contact disk lies on a *radical plane* defined by an intersection of two spheres, and intersections of radical planes are known to define the *radical Voronoi tessellation* (also known as the *Laguerre-Voronoi tessellation* (Imai et al., 1985) or the *power diagram* (Aurenhammer, 1987)). Therefore, the intersections of a contact disk with other contact disks can produce a portion of a radical Voronoi cell face that is constrained inside the initial contact disk. Such a constrained contact surface respects the geometric partitioning of space between the atomic balls. It also implicitly considers the presence of solvent by not extending outside of the solvent-accessible surface of a molecular structure. Thus, we assume it is a valid representation of an interatomic interaction.

### 2.2. Atom-atom contact construction algorithm

Given a pair of atoms *a* and *b* that have a contact disk, without loss of generality, let us assume that *a* has a smaller set of close-enough neighbors (for example, the upper green ball in Fig. 1a and 1b). This set contains all the possible balls that are close-enough Voronoi neighbors to both atoms. We set the initial contact representation to be the *full contact disk* between *a* and *b*. We then iteratively cut it with every contact disk defined for *a* and its close-enough neighbors. Figure 1c illustrates this process. After every cut, we are left with the portion of the contact surface that lies in the same half-space of the cutting disk plane as *a*. We stop the process when there is no surface left (meaning that *a* and *b* are not in contact anymore), or after all the neighbors of *a* were processed.

While our algorithm naively follows the definition of the radical Voronoi tessellation without employing any clever heuristics, there are several aspects that make it practical.

Firstly, in molecular structures, the density of the atomic packing tends to be not too far from the level allowed by the laws of physics. Therefore, the number of close-enough neighbors for every atom are bound by a constant value when using a solvent probe radius close to 1.4 Å. This allows employing spatial hashing using a regular grid of buckets to efficiently query all the relevant neighbors for every atom.

Secondly, we employ only computationally cheap geometric operations when performing intersection filtering and cutting operations. In the iterative portion of the algorithm, we initially represent the contact disk by its minimal surrounding hexagonal polygon, we cut that polygon iteratively, and only when the cutting is over we replace polygon line segments with circular arcs where needed.

Thirdly, different contacts are constructed independently from each other, and the final result does not depend on the order of the construction. Therefore, the algorithm is trivially parallelizable. Moreover, the algorithm is not required to construct all contacts if we need just a subset of them. For example, we can choose to compute only inter-residue or inter-chain contacts.

### 2.3. Contacts updating algorithm

Consider a scenario where we have a set of atomic balls with an associated set of constructed contacts, and we need to modify (move or resize) a subset of these balls and subsequently update the contacts. One possible motivation for such a scenario is to incorporate contacts-based analysis into molecular simulations — for example, by extending modular frameworks for Monte Carlo simulations such as Faunus (Stenqvist et al., 2013). Based on our definition of contacts, the modification of the balls can affect the following categories of contacts: the contacts between the modified balls; the contacts between the modified balls and their close-enough neighbors (close-enough before or after the modification); the contacts between the close-enough neighbors of the modified balls. Any other contacts cannot be affected because the modified balls cannot interfere with them. Let us denote the set of indices of the modified balls as *I*_*m*_. The modified balls can have different sets of close-enough neighbors before and after the modification: let 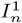 represent the indices of close-enough neighbors before the modification and 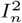 the indices of close-enough neighbors after the modification. The overall set of indices of the balls relevant to updating the contacts is 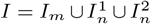 This allows us to define the contact update procedure in Voronota-LT as a simple two-step process: before modifying the balls, delete all the contacts between any pair of balls whose indices are both in *I*; after the modification of balls, reconstruct all possible contacts between any pair of balls whose indices are both in *I*.

### 2.4. Periodic boundary conditions application algorithm

Representing molecular systems as periodic lattices of identical subunits (*unit cells*) is common when performing molecular dynamics or Monte Carlo simulations (Frenkel and Smit, 2002). It is also crucial for the analysis of molecular crystals, which is commonly carried out using crystal (or quotient) graphs (Xie and Grossman, 2018) that account for both the intra- and inter-unit cell interactions. Consider a scenario when we have to construct interatomic contacts for a molecular system in a periodic lattice that is defined by three translation vectors. We can use those vectors to generate a *supercell* composed of 3 × 3 × 3 = 27 unit cells, providing a complete environment for the unit cell in the center of the 3 × 3 × 3 grid. We then construct only the contacts that involve at least one atom from the central unit cell. Contacts between atoms from different unit cells are repetitive — there can be from two to six symmetrical copies of every such contact, but we only construct a single copy to calculate and report the contact area.

### 2.5. Constrained Voronoi cell characterization algorithm

We define the *constrained Voronoi cell* of an atom as the space region bounded by the contact surfaces and the contact sphere of that atom (Fig. 1d). Such a cell may be not fully closed because some contacts can have circular arc edges. Such arcs lie on the contact sphere surface and emborder regions of the solvent-accessible surface of the atom (colored in red in Fig. 1d), that is, parts of the contact sphere where a solvent probe center can be placed without being inside any neighboring Voronoi cell.

To compute the total solvent-accessible area corresponding to the constrained Voronoi cell, we use the fact that the area of a region on a sphere surface can be directly calculated from the solid angle subtended by the region from the sphere center (Todhunter, 1863). Instead of computing the solid angle of the solvent-accessible surface directly, we calculate the solid angles for all the atom-atom contact surfaces, sum them up, and subtract the sum from the total solid angle of a sphere that equals 4*π* steradians. As every our contact is a convex polygon with sides that are either line segments or circular arcs, we can compute every contact-subtended solid angle analytically. There is a pitfall in the radical Voronoi tessellation when the sphere’s center lies outside the sphere’s Voronoi cell, which can make the solid angles subtended by the cell faces from the sphere’s center overlap. We handle such overlaps by inverting solid angle values (subtracting them from 4*π*) where needed.

Knowing the total solid angle of the solvent-accessible regions also allows us to compute the volume of the corresponding spherical sector inside the contact sphere. To get the total volume of the constrained Voronoi cell, we add the total sum of volumes of the pyramids formed by using the sphere’s center as the apex and the contact surfaces as bases. In cases when the sphere’s center lies outside the Voronoi cell, the pyramids can overlap. Therefore, we invert pyramid volumes (multiply them by −1) where needed. We characterize Voronoi cells using only contact polygons without the need of explicitly pre-computed Voronoi vertices or edges. It makes our approach robust in situations where input contains arrangements of balls that induce degenerate Voronoi vertices or edges, that is, vertices shared by more than four cells or edges shared by more than three cells (Fig. 1efg). Such situations can frequently occur in molecular structures as they often contain symmetrical parts.

### 2.6. Implementation

We implemented the contact construction and cell characterization algorithms in C++ as a header-only library and as a standalone command-line tool, both called “Voronota-LT”. The implementation includes the contacts and cells updating and the periodic boundary conditions handling features. Given an input list of atomic balls with labels that include chain and/or residue identifiers, there are options to limit Voronota-LT to compute only inter-residue or inter-chain contacts. The software also provides access to the coordinates of the constrained Voronoi face vertices that can be used for 3D visualization of contacts. The parallelization is implemented using OpenMP (Dagum and Menon, 1998) and can be enabled during compilation. Apart from the optional usage of OpenMP, both the library and the command-line tool do not depend on anything but the C++ standard library.

### 2.7. Comparison of algorithms in Voronota-LT and in other methods

Compared to Voronota-LT, the vanilla Voronota software (Olechnovič and Venclovas, 2014, 2021) employs a substantially different approach to constructing atom-atom contacts. Given an input set of balls, Voronota first computes all the vertices of the additively weighted Voronoi tessellation, and then uses the vertices to identify neighbors that are both close enough and have adjacent Voronoi cells. Then, every contact is constructed starting from the initial triangulated representation that corresponds to the intersection of a contact sphere with a hyperboloidal surface (which is a plane when the two contacting atoms have the same radius). The obtained triangulated representation is then iteratively updated by cutting it with neighboring hyperboloidal surfaces. The resulting contact surface can be non-convex and can have disconnected parts. The solvent-accessible surface is constructed from the triangulated representation of a contact sphere cut by neighboring contact spheres.

In contrast, Voronota-LT does not pre-compute Voronoi vertices, uses a much simpler representation of a contact surface, and calculates all the areas and volumes analytically.

Additionally, the general Voronota-LT methodology for constructing contacts allows for the usage of different forms of the Voronoi diagram. Therefore, we could additionally implement the construction of constrained triangulated contact surfaces that correspond to the additively-weighted Voronoi tessellation, like in vanilla Voronota. We called this variation “Voronota-LT-AW” (“AW” stands for “Additively Weighted”), while the default, simpler and faster variation that uses the radical Voronoi tessellation is called “Voronota-LT-Radical” or just “Voronota-LT”.

When comparing Voronota-LT to methods that compute tessellation-based contact areas as an integral part of calculating atomic volumes, namely ALPHAVOL (Edelsbrunner and Koehl, 2003), POWERSASA (Klenin et al., 2011) or SBL-Vorlume (Cazals et al., 2011), the key distinction is that Voronota-LT does not derive contacts from the fully constructed Voronoi diagram or its dual triangulation. Because Voronota-LT does not rely on the full Voronoi tessellation, it is especially efficient when exclusively calculating specific subsets of contact areas, for example, protein-protein interaction interfaces. Moreover, Voronota-LT naturally supports parallel contact computation. Its contact-level approach to parallelization differs from the Voronoi cell-level parallelization employed by methods such as Voro++ (Lu et al., 2023). When focusing on specific interfaces, there is no need to compute all contacts for every involved Voronoi cell. Therefore, contact-level parallelization is likely to be more efficient for interface studies.

## 3. Benchmarking

### 3.1. Data

We benchmarked the Voronota-LT command-line tool on a diverse set of structures of multi-chain biological assemblies downloaded from the Protein Data Bank (PDB) (wwPDB consortium, 2019). We queried the PDB using RCSB.org GraphQL-based API (Rose et al., 2021) for protein and nucleic acid entities that belonged to the PDB entries deposited before 2024-01-03 and where the first biological assembly contained two or more polymer chain instances (with up to 5000 polymeric residues overall) and at least one non-polymer entity instance. We took the PDB IDs of the entities that were cluster representatives at 30% sequences identity. The full GraphQL query is given in Fig. S1. For every PDB ID, we downloaded the first biological assembly in the mmCIF format and converted it to a labeled list of atomic balls with the Voronota software, taking only heavy (non-hydrogen) atoms. The van der Waals radii were assigned using the previously published atom type-to-radius mapping (Li and Nussinov, 1998; Olechnovič and Venclovas, 2021). We then filtered out structures with extreme clashes where two or more balls had exactly the same center. In total, we obtained 14,861 oligomeric molecular structures with atom counts ranging from 256 to 151,585.

### 3.2. Comparative analysis of interatomic contacts computation

For every prepared input structure, we computed atom-atom contact areas with the Voronota-LT method in five different modes: “LT-Radical” (default mode, computing all atom-atom contacts areas and also computing all solvent-accessible surface areas), “LT-Radical parallel” using 10 OpenMP threads (computing the same areas as in the default mode), “LT-Radical inter-residue” (computing only contact areas between atoms from different residues), “LT-Radical inter-chain” (computing only contact areas between atoms from different chains), “LT-AW” (computing atom-atom contact areas that correspond to the additively-weighted Voronoi tessellation). We also ran vanilla Voronota software in its default mode (denoted as “Vanilla-AW”) that computed atom-atom contacts and solvent-accessible surface areas. All the software was compiled and run on the same machine (20-core Intel® Xeon® Silver 4210R CPU @ 2.40GHz, Ubuntu Linux 23.04, GCC g++ compiler version 12.3.0).

We recorded the wall running time values and plotted them in Fig. 2. On average, when compared to vanilla Voronota, “LT-AW” is only twice faster, whereas “LT-Radical” is 16 times faster. The Voronota-LT-Radical modes that only compute inter-chain and inter-residue contacts are, respectively, 23 and 105 times faster than “Vanilla-AW”. Moreover, as evident from the middle and the right parts of Fig. 2 that show running times per 10,000 input atoms, the efficiency of Voronota-LT (in any mode) does not decrease when the input size increases.

**Fig. 2:**
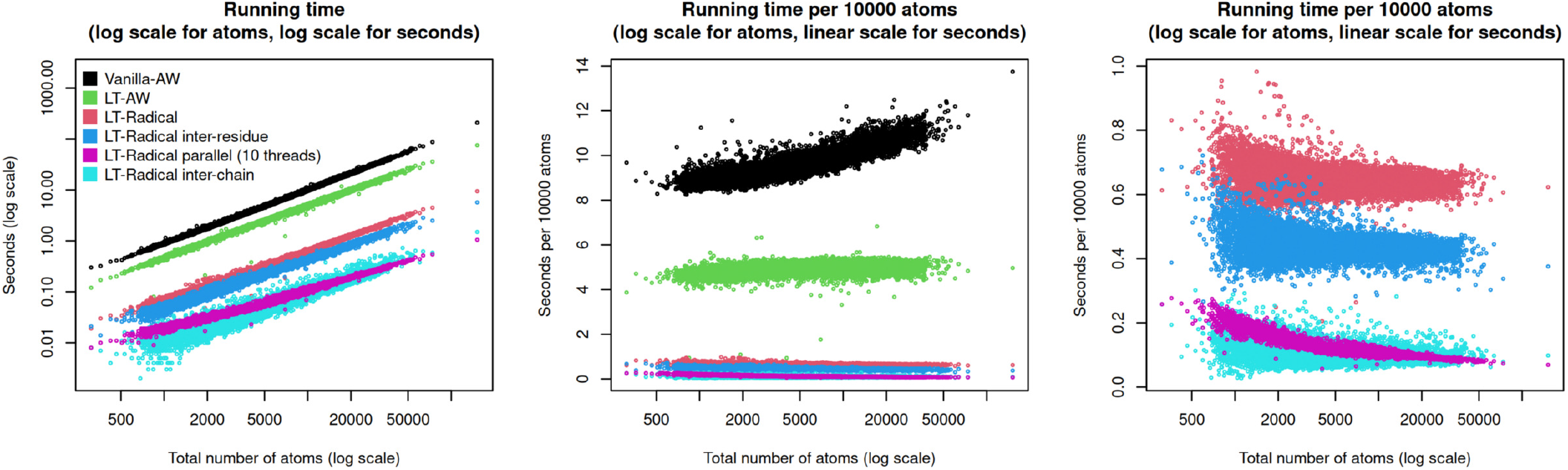
Time efficiency of constructing contacts using vanilla Voronota (Vanilla-AW) and various modes of Voronota-LT (LT-*). Total wall running times were used for every plot.

We also computed the Pearson correlation coefficients between differently produced sets of areas for every input structure (Table S1). We summarized every distribution of coefficients by calculating the minimal (*ρ*_min_) and the average (*ρ*_avg_) values. When all the input ball radii were set to 1.8 Å, the vanilla Voronota and Voronota-LT contact areas correlated ideally (*ρ*_min_ = 1, *ρ*_avg_ = 1), as expected. When the radii were not changed to be the same, the vanilla Voronota and Voronota-LT-AW contact areas correlated almost ideally (*ρ*_min_ = 0.999, *ρ*_avg_ = 1) — we attribute the slight differences to the usage of triangulated approximations of surfaces for computing areas.

Because the differences between van der Waals radii of different atoms in biological macromolecular structures are not extreme, contact areas based on the radical and additively weighted Voronoi tessellations were expected to correlate highly. Indeed, vanilla Voronota and Voronota-LT-Radical contact areas correlated with *ρ*_min_ = 0.9609 and *ρ*_avg_ = 0.976, the histogram of coefficients is presented in Fig. S2. Interestingly, the contact areas summarized on the residue-residue level correlated even higher (*ρ*_min_ = 0.9947, *ρ*_avg_ = 0.999). The summarized chain-chain areas correlated even better (Fig. S3), the Pearson correlation coefficient calculated for all 67,700 interfaces in the benchmark dataset was greater than 0.9999. Solvent-accessible surface areas computed by vanilla Voronota and Voronota-LT also correlated almost ideally (*ρ*_min_ = 0.9997, *ρ*_avg_ = 1).

### 3.3. Comparative analysis of atomic volumes computation

We compared the performance of Voronota-LT in computing per-atom volumes with the performance of SBL-Vorlume tool (Cazals et al., 2011) from the Structural Bioinformatics Library (Cazals and Dreyfus, 2017). We used 12,403 PDB structures for which we were able to ensure that Voronota-LT and SBL-Vorlume read exactly the same set of atoms. We used atomic radii assigned by SBL-Vorlume and set the rolling probe radius to 1.4 Å. We executed SBL-Vorlume in two modes: the default (faster) and the exact (more precise and robust, but slower). We executed Voronota-LT without parallelization. We recorded the calculation times reported by the software tools, the time spent on input/output was not considered. The results are presented in Fig. 3a, Voronota-LT was at least several times faster in every case.

**Fig. 3:**
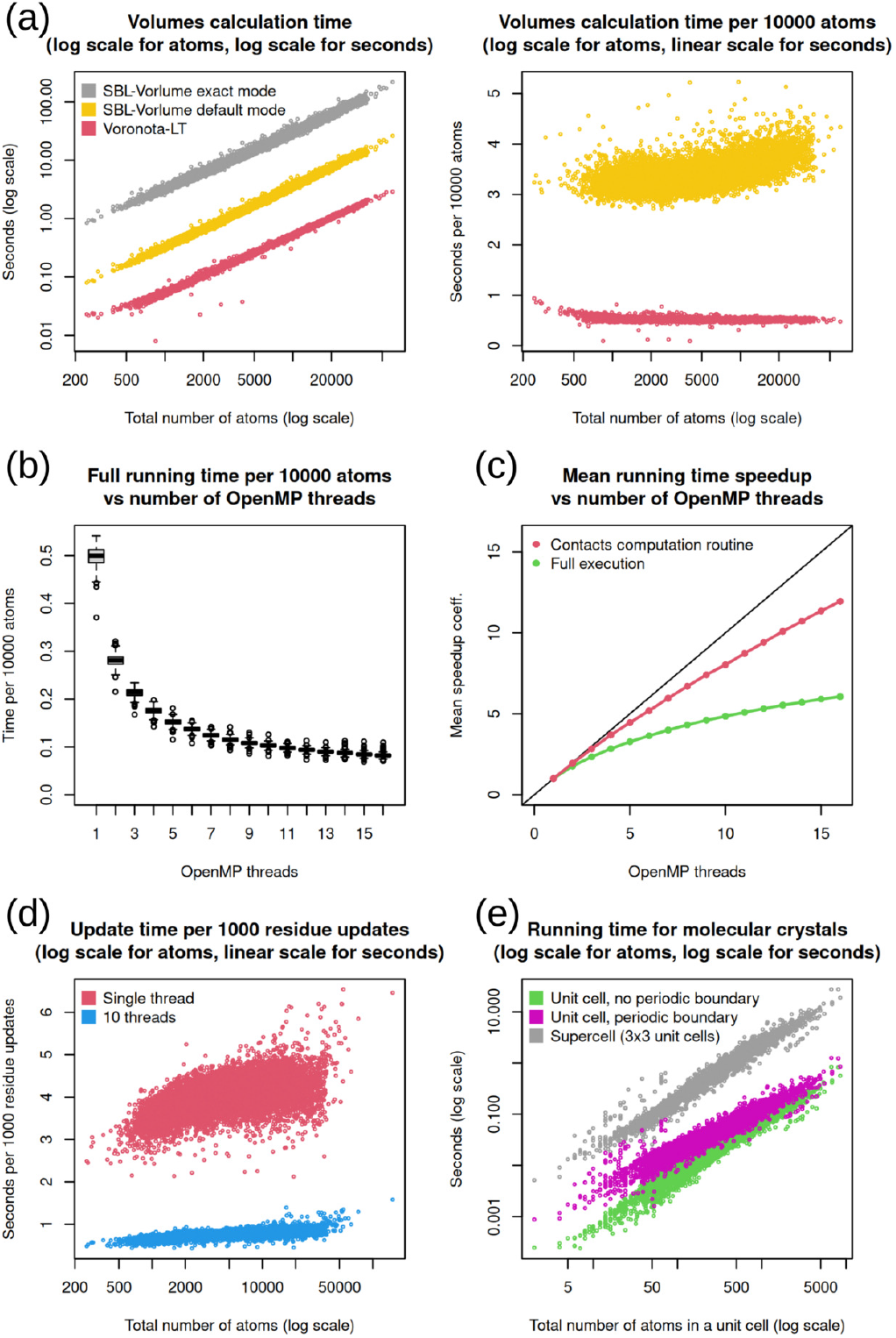
**(a)** Comparative performance of atomic volumes computation using Voronota-LT and SBL-Vorlume. Reported calculation times were used, input/output times were not considered. **(b)** Voronota-LT full running time distributions for the number of OpenMP threads ranging from 1 to 16. **(c)** Parallel speedup of Voronota-LT computations. **(d)** Performance of the Voronota-LT update procedure. **(e)** Performance of Voronota-LT when processing molecular crystal structures.

We also compared the calculated atomic volumes — they were very similar. Indeed, for most structures, the maximum observed difference between volumes of the same atoms computed using different methods was less than 0.001, and such difference can often be attributed to different settings of floating point input/output. Interestingly, the results of Voronota-LT were more similar to the results of the exact mode of SBL-Vorlume (see Fig. S4). We consider SBL-Vorlume to be a reliable benchmark because its implementation is based on the state-of-the-art Computational Geometry Algorithms Library (CGAL, https://www.cgal.org). In Voronota-LT, volumes are calculated from the contact-derived solid angles, thus the correctness of volumes also serves as an experimental evidence of the correctness of contacts.

### 3.4. Parallelization efficiency analysis

The Voronota-LT-Radical executed in parallel using 10 threads performed, on average, 6.3 times faster than the unparallelized Voronota-LT-Radical (Fig. 2a). This is not surprising because only the contacts computation procedure is effectively parallelized in Voronota-LT. To further investigate the parallelization efficiency, we took the 500 largest input structures (with atom counts ranging from 22,169 to 151,585), and processed them with Voronota-LT-Radical using different numbers of OpenMP threads. We calculated the average parallel speedup coefficients for every number of threads, for both the total running times and the contacts computation procedure running times. The plot in Fig. 3b shows what improvements of time per 10,000 atoms to expect when using different numbers of threads. The speedup plot in Fig. 3c shows that the computation of the contacts is parallelized relatively efficiently, especially for a lower number of threads where the speedup is close to ideal.

### 3.5. Performance of contacts updating

We used the previously introduced set of 14,861 PDB structures to benchmark the average time it takes Voronota-LT to update the set of contacts and the set of atomic cell descriptors after moving atoms of a single residue (in the analyzed set, on average, one residue has 7.9 atoms). For every input structure, we repeated 500 times the following iteration that involved two updates: moved a randomly selected residue in a randomly sampled direction by a distance randomly sampled from the [1.0; 5.0] Å interval; updated the contacts and the cells; moved the perturbed residue back; updated the contacts and the cells. We recorded the calculation times and confirmed that there were no accumulated errors — the contacts and the cells before and after all the perturbing/restoring modifications were identical. The recorded times are presented in Fig. 3d. Using a single thread it usually takes 3-5 seconds to perform 1000 residue updates. However, the updating procedure benefits from parallelization — using 10 OpenMP threads, 1000 residue updates usually take less than one second.

### 3.6. Performance with periodic boundary conditions

We tested Voronota-LT on crystal structures from the Crystallography Open Database (COD) (Gražulis et al., 2009). Using the OPTIMADE API (Andersen et al., 2021), we downloaded sets of unit cell atom coordinates and the corresponding lattice vectors for 10,894 COD entries published in 2023. We processed every crystal structure in three regimes: as an unbound isolated unit cell without any boundary conditions (to obtain the minimal processing time for a unit cell); as a unit cell with periodic boundary conditions applied (see Fig. S5 for examples of constructed contacts); as a supercell formed from a 3 × 3 × 3 grid of unit cells. Fig. 3e shows that the handling of periodic boundary conditions in Voronota-LT did not substantially slow down the processing of the unit cell, especially for larger unit cells with over 500 atoms. In any case, handling periodic boundary conditions was markedly more efficient than the supercell processing.

## 4. Application examples

### 4.1. Studying interactions in protein-protein interfaces

The Voronoi cell of a single atom *a* fully defines the neighborhood of *a* in terms of the closest space points, while the corresponding Voronoi faces describe how the other atoms influence *a* by having common closest space points, and that influence can be quantified by constructing tessellation-based contacts and calculating their areas. Those contacts can be grouped to describe different categories of interactions — for example, inter-chain interactions in protein complexes, as showcased in Fig. 4a.

**Fig. 4:**
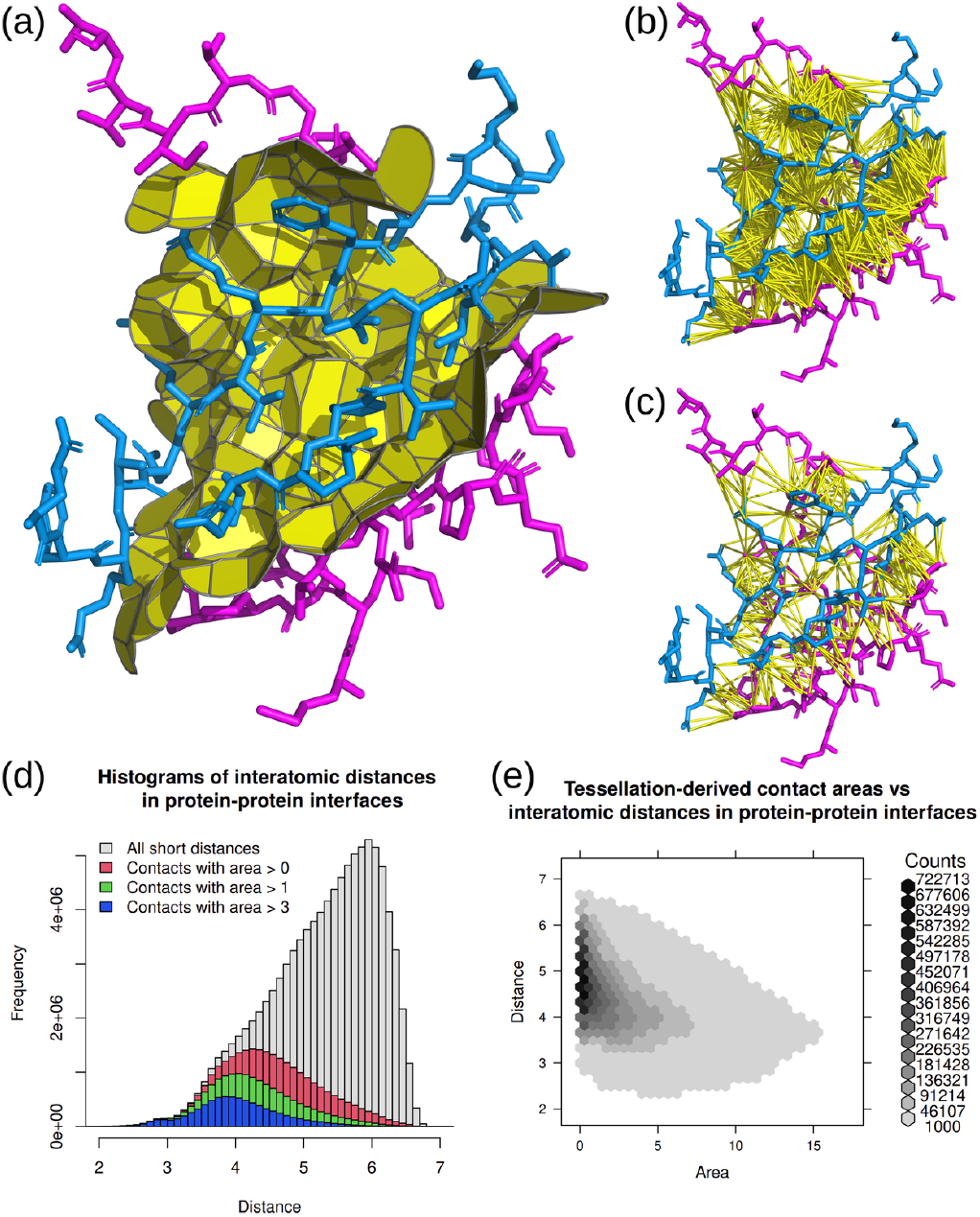
**(a)** Tessellation-derived inter-chain interface contacts, calculated for the first model from the PDB entry 1CIR. **(b)** Distance cutoff-derived pairs of atoms in an inter-chain interface. **(c)** Pairs of atoms in an inter-chain interface that have tessellation-derived contacts. **(d)** Histograms of distances between atoms in inter-chain interfaces observed in the processed PDB structures. Four overlapping categories of interactions are analyzed: interactions that satisfy the distance cutoff (gray), interactions that have tessellation-derived contacts with any area (red), with area greater than 1Å^2^ (green), or with area greater than 3Å^2^ (blue). **(e)** Two-dimensional hexagonal histogram of areas and distances of tessellation-derived contacts in inter-chain interfaces observed in the processed PDB structures.

Based on our definition of contacts, we consider interactions between atoms that are close-enough, meaning that a water-sized probe (1.4Å radius ball) cannot fit between them. If we only consider this distance cutoff-based condition, the number of inter-chain atom pairs can be overwhelming even for a relatively small interface (Fig. 4b). Only a relatively small fraction of those pairs involve atoms that share a tessellation-based contact (Fig. 4a,c). In such pairs, the atoms have at least some closest space points in common, therefore such pairs are more relevant for describing the inter-chain interface.

Let us investigate the relationship between the distance cutoff-derived and tessellation-derived interactions quantitatively, using the previously introduced set of 14,861 oligomeric molecular structures from PDB. Fig. 4d shows the histogram (in gray) of the distances between the centers of atoms for all the 98,768,354 pairs of atoms that are close-enough to have an inter-chain contact according to the Voronota-LT definition. The histogram in red summarizes the distances between the centers of atoms that are actually in contact, that is, have a non-zero contact area calculated by Voronota-LT. Among all the observed close-enough pairs of atoms, less than 27% form a tessellation-based contact. Contacts with larger areas are more rare — for example, the contacts with area greater than 3Å^2^ correspond to less than 8% of all the close-enough pairs (see the blue histogram in Fig. 4d).

Because the area of a contact depends not only on the distance between the corresponding interacting atoms but also on how the contact is influenced by the surrounding atoms, we expect that there is no trivial relationship between the interatomic distances and the areas of tessellation-derived contacts. Our data analysis results support this expectation. According to Fig. 4d, a greater distance does not necessarily mean absence of a contact area. Also, when we exclusively consider the tessellation-derived contacts, there is no high correlation between areas and distances (Pearson correlation coefficient ≈ −0.43, see Fig. 4e for the two-dimensional histogram plot).

This means that tessellation-based contact areas is valuable information that cannot be easily derived from ordinary distance-based analysis. One of possible uses of such information is studying interactions in structural ensembles. Calculating simple statistics of distributions of contact areas, for example, variance values, can help to characterize the variability of every interaction without disregarding the effects of its structural environment. Fig. S6 presents a toy example of a simple contact area variability analysis, where we analysed the NMR ensemble from the study on chymotrypsin inhibitor 2 folding (Neira et al., 1996) and visualized the relative variances of contact areas on both atom-atom contact and the residue-residue levels. Thanks to the speed of Voronota-LT, this type of analysis can be performed for much larger molecules with many more structural states. Another benefit of the tessellation-based variability analysis is that structural states do not need to be superposed, that is, structurally aligned.

### 4.2. Studying contact changes in large structures from time-resolved cryo-EM experiments

Modern time-resolved cryo-EM experiments can provide structural models of big molecular complexes at various stages of biological processes. Analysis of such data can be challenging due to its scale and complexity. We present an example of how a tessellation-based analysis can help identify and describe changes in interactions between chains in a 70S *E. coli* ribosome that undergoes splitting of its 50S and 30S subunits mediated by an external protein factor. We took structural data from the time-resolved cryo-EM study of HflX-mediated ribosome recycling (Bhattacharjee et al., 2023). We analysed differences between inter-chain interface contacts computed for two ribosome structures determined on an earlier (PDB ID 8G34) and on a later (PDB ID 8G38) stages of splitting (Fig. 5a). We computed inter-chain contact areas on the residue-residue level and calculated contact area differences between different structural states. Inter-chain interfaces inside the 50S and 30S subunits did not differ systematically, but the contact changes in the interfaces between 50S and 30S were dramatic (Fig. 5b). The tessellation-based analysis allowed us to describe the almost complete rearrangement of the contacts between the 50S and 30S subunits (Fig. 5c), and the emergence of an additional interface involving the HflX factor in the later stage of the splitting (Fig. 5d).

**Fig. 5:**
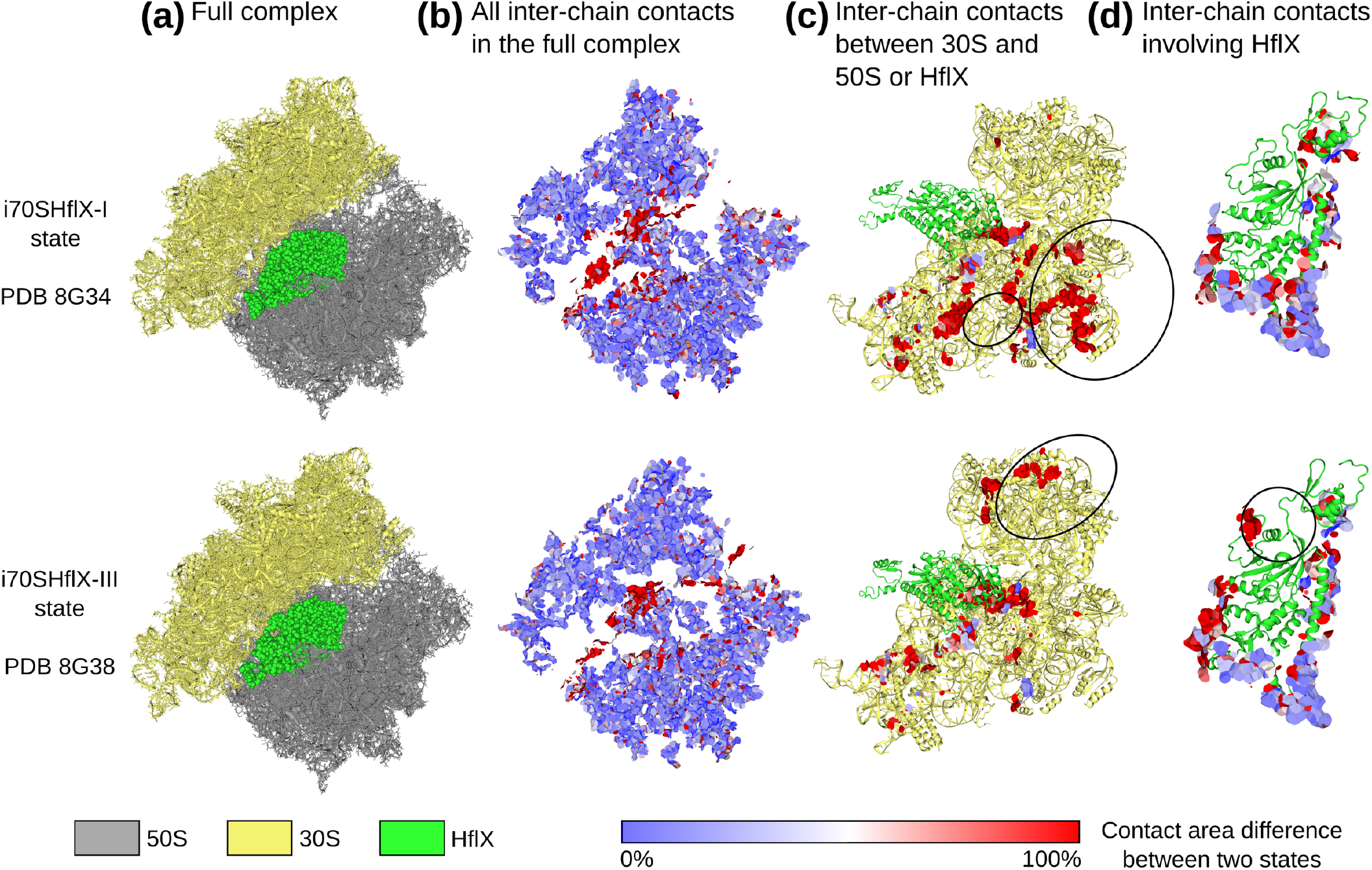
Example application of Voronota-LT for studying inter-chain interface contact differences between two structural states of the 70S ribosome in complex with HflX protein that acts as a splitting factor between 50S and 30S subunits. The analysed structures were taken from the Protein Data Bank entries 8G34 and 8G38 deposited by the authors of the time-resolved cryo-EM study of HflX-mediated ribosome recycling (Bhattacharjee et al., 2023). 3D visualizations were generated using Voronota-GL.

For another example, shown in Fig. S7, we computed inter-chain interface contact areas for the contracted (PDB ID 8C38) and the extended (PDB ID 8CPY) structural states of the cowpea chlorotic mottle virus capsid determined during the time-resolved cryo-EM study of viral dynamics (Harder et al., 2023). Each structure had 180 protein chains of the same sequence. We evaluated the differences in chain packing: the contracted variant had 786 inter-chain interfaces with a total area of 365,093 Å^2^, while the extended variant had 390 inter-chain interfaces with a total area of 184,506 Å^2^.

## 5. Discussion and conclusions

We presented Voronota-LT, a method tailored to a very specific task — computing and reporting tessellation-based atom-atom contacts constrained inside the solvent-accessible surface of a 3D molecular structure. Unlike other tessellation-inspired methods, Voronota-LT constructs every contact directly, without needing any pre-computed global Voronoi diagram or Delaunay triangulation. This allows efficient focusing on subsets of interactions, for example, inter-chain interfaces.

Voronota-LT has some limitations. It will not work efficiently for input sets of balls that are not similar to molecules, for example, when balls are much more densely packed or have vastly different radii, or when using a solvent probe radius much larger than 1.4 Å. However, when used for the problem that it was designed for, Voronota-LT is, to the best of our knowledge, the most efficient tool available.

We evaluated Voronota-LT on a dataset of 14,861 protein complex structures containing approximately 90 million heavy (non-hydrogen) atoms in total. Processing this dataset, including reading the input and computing atom-atom contact areas, solvent-accessible areas, and atomic volumes, took about 96 minutes on a single CPU core. As shown in Fig. 2, empirical analysis indicates that Voronota-LT’s runtime scales linearly with the number of input atoms. Using this scalability, we can estimate the time required to process a large protein structure database, such as AlphaFold Protein Structure Database (AlphaFoldDB) (Varadi et al., 2022). AlphaFoldDB consists of models of proteins from the UniProtKB/TrEMBL database (UniProt Consortium, 2023) that, as of 2024-10-02, contains over 248 million protein sequence entries, comprising over 88 billion amino acids. Based on the amino acid frequences in UniProtKB/TrEMBL, AlphaFoldDB is estimated to contain approximately 684 billion heavy atoms. With access to a cluster of 1,024 CPU cores, Voronota-LT could process all AlphaFoldDB models in about 12 hours. On a single workstation with 40 CPU cores, the task could be completed in about 13 days. While these estimations are approximate, they highlight the practicality of using Voronota-LT for large-scale analysis, even without access to expensive computational resources.

Most importantly, the efficiency of Voronota-LT is coupled with the ability of tessellation-derived contacts to describe interactions in a comprehensive manner, fully considering the relevant structural context and providing meaningful and summarizable interaction value descriptors — contact areas. Currently, there are multiple methods that use vanilla Voronota contacts, for example: CAD-score, VoroMQA, VoroIF-GNN methods used in the VoroIF-jury pipeline (Olechnovič et al., 2023); VoroCNN (Igashov et al., 2021a); S-GCN (Igashov et al., 2021b). In principle, all those methods can potentially use Voronota-LT contacts and run significantly faster. We provide Voronota-LT freely with a permissive open-source code license to enable the broad structural bioinformatics community to benefit from the tessellation-based analysis of molecular interactions without much computational resources.

## Supplementary information

**Fig. S1.**
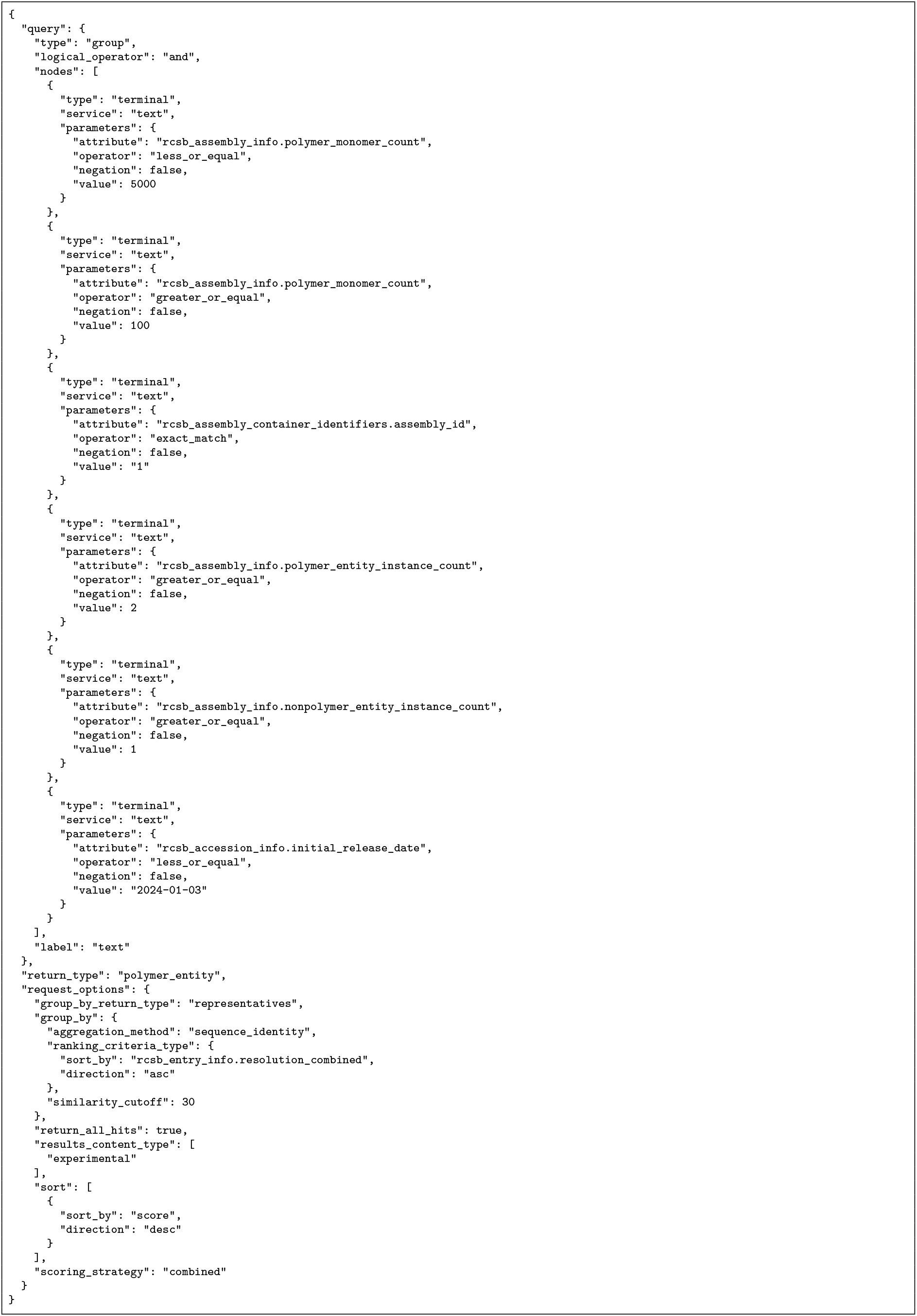
RCSB GraphQL query used to download a list of PDB entities to get PDB entry identifiers for the benchmarking of Voronota-LT.

**Fig. S2.**
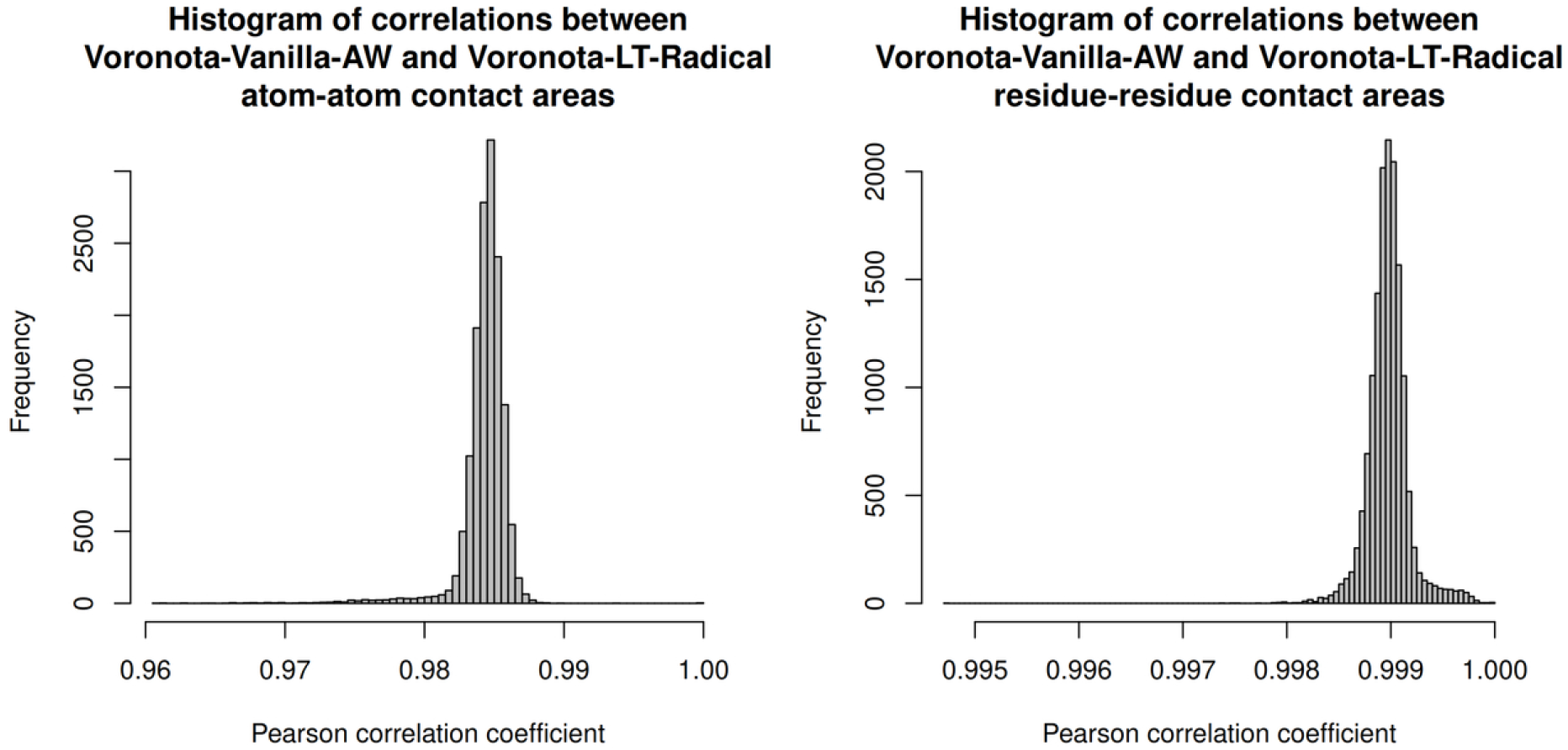
Correlation between contact areas computed by vanilla Voronota and Voronota-LT on atom-atom and residue-residue levels.

**Fig. S3.**
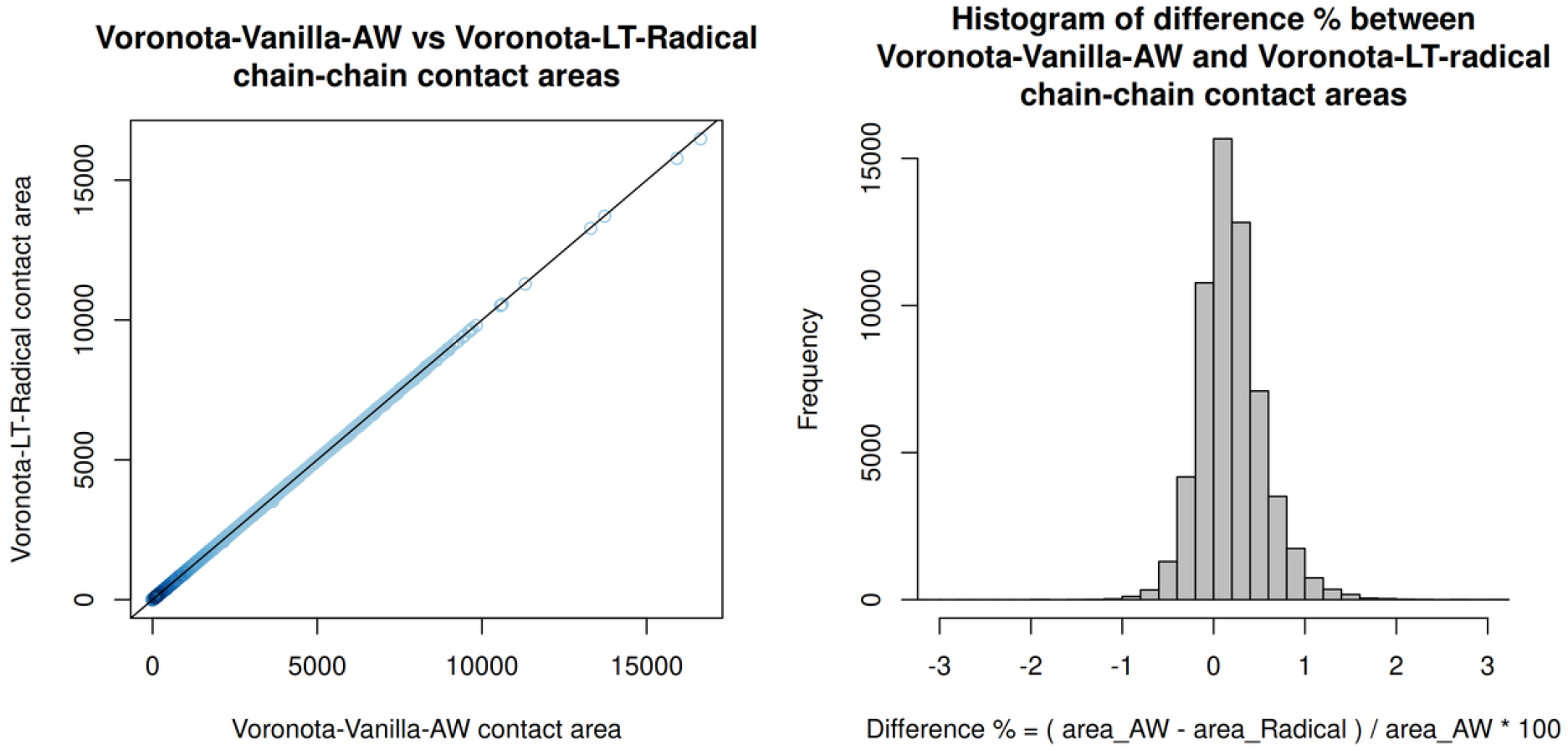
Correlation between inter-chain interface contact areas computed by vanilla Voronota and Voronota-LT.

**Fig. S4.**
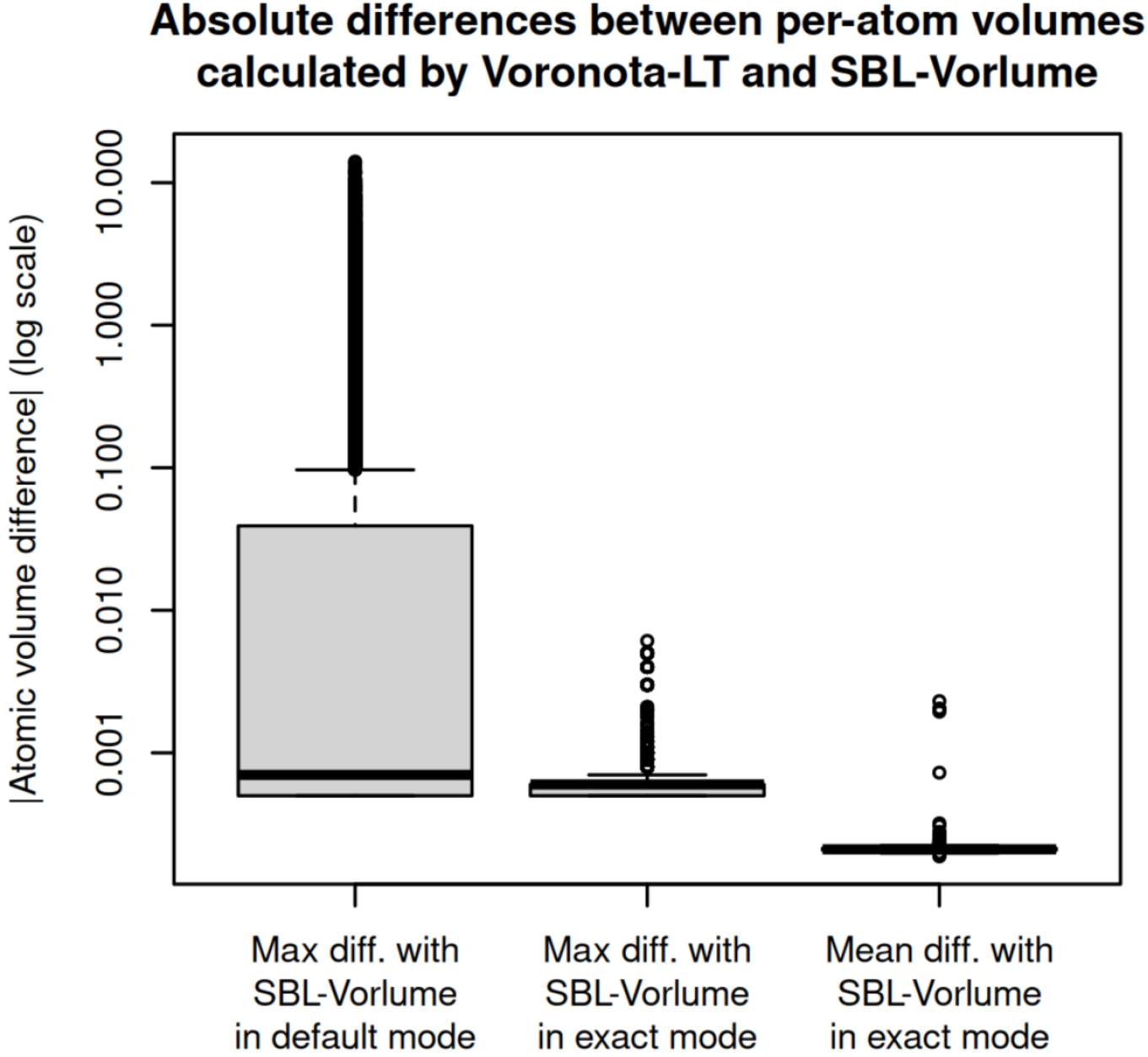
Distributions of maximum and mean differences of per-atom volumes calculated by Voronota-LT and SBL-Vorlume for the analyzed protein structures.

**Fig. S5.**
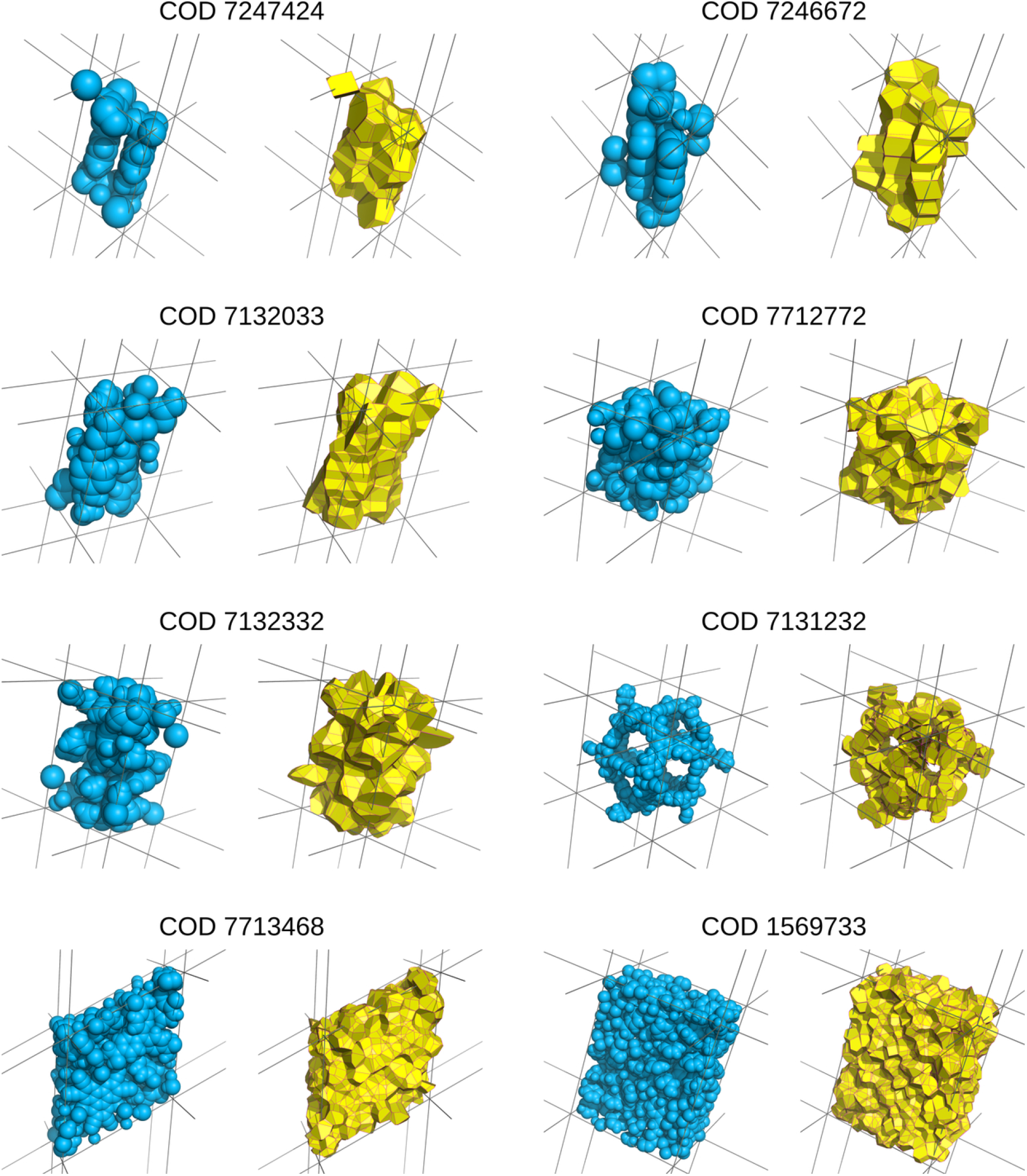
Examples of the Voronoi-Laguerre tessellation-derived contacts computed for several crystals from the Crystallography Open Database (COD). The contacts (colored in yellow) were computed for the unit cell atomic balls (colored in blue) considering the periodic boundary conditiones defined by the cryslall lattice vectors (indicated using the gray lines). The relevant COD entry identifiers are displayed above every example.

**Fig. S6.**
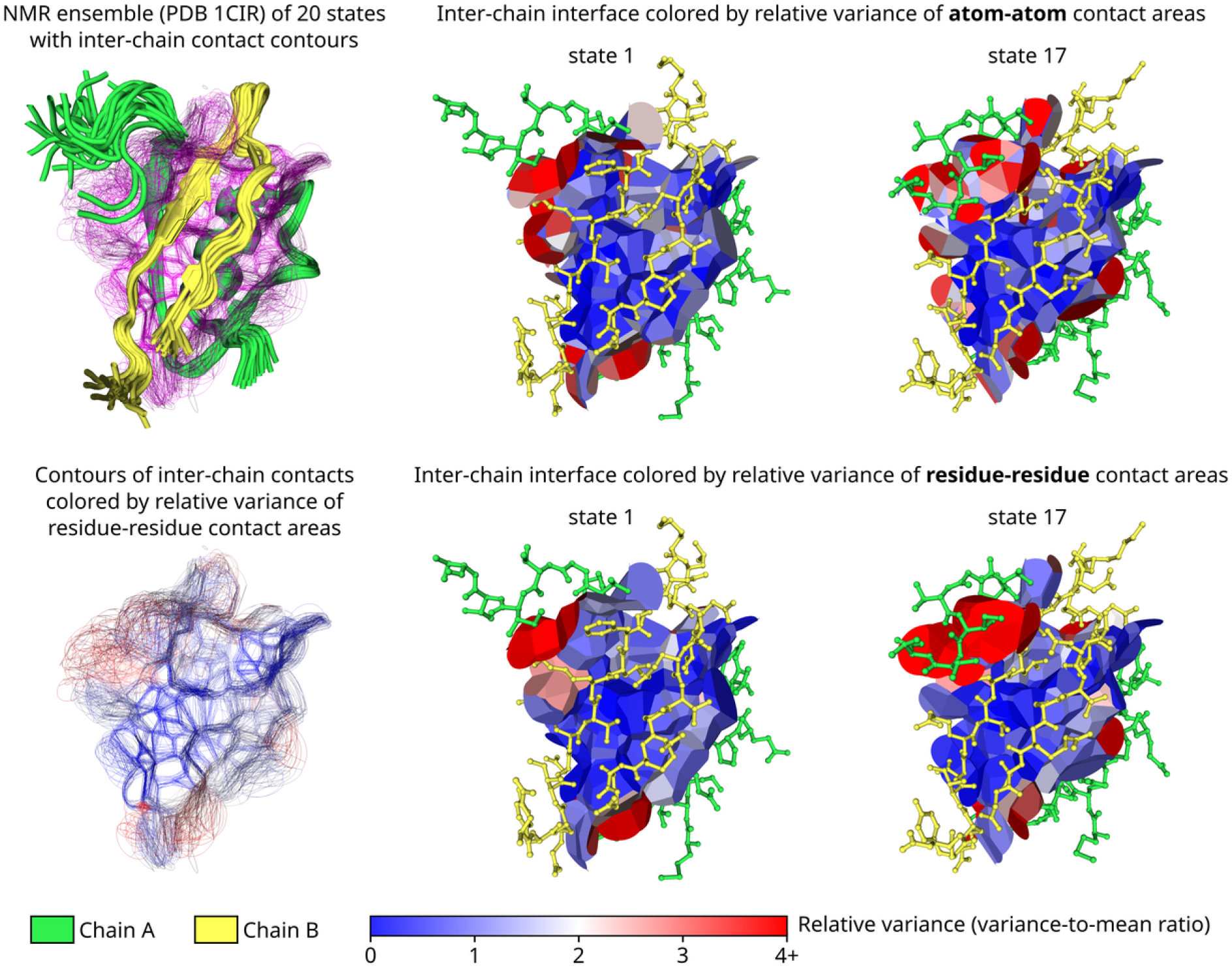
Example application of Voronota-LT for studying inter-chain interface contact variability in an ensemble of dimeric protein structures. We analysed an NMR ensemble of 20 conformational states taken from the Protein Data Bank entry 1CIR deposited by the authors of the study on chymotrypsin inhibitor 2 folding (Neira et al., 1996). It is a dimeric protein molecule, and we focused on the contacts in the inter-chain interface. We computed the contact areas and, for every unique contact, we calculated the mean and the variance values of the corresponding observed area distribution. We considered both atom-atom contact areas and the residue-residue level areas derived by summing atom-atom areas according to residue identifiers. We visualized the results by displaying inter-chain contact surfaces and coloring them by the corresponding variance-to-mean ratios (relative variances) that indicate how much every contact area is dispersed relative to its mean. We observed what contact areas varied the least and the most. In particular, the contact areas further from the solvent, in the interface core, were more stable. The most variable interface region involved interactions with a C-terminal tail of chain A. 3D visualizations were generated using Voronota-GL.

**Fig. S7.**
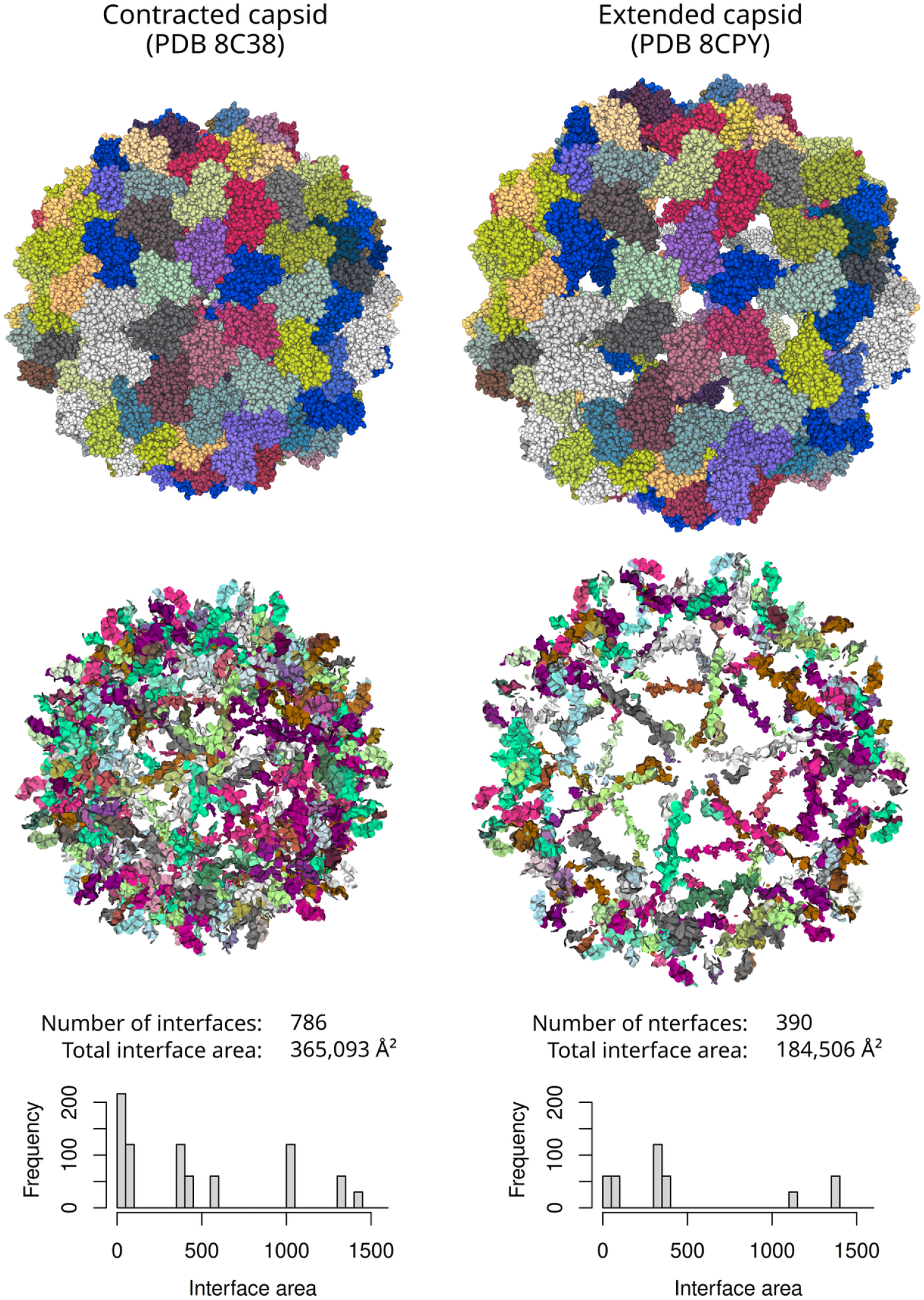
Example application of Voronota-LT for studying inter-chain interfaces in contracted and expanded structures of the cowpea chlorotic mottle virus capsid. The analyzed structures were taken from the Protein Data Bank entries 8C38 and 8CPY deposited by the authors of the time-resolved cryo-EM study of viral dynamics (Harder et al., 2023). Each structure had 180 protein chains of the same sequence, the total number of atoms was 215,580 for 8C38 and 198,420 for 8CPY. The contracted variant had 786 inter-chain interfaces with a total area of 365,093 Å^2^, while the extended variant had 390 inter-chain interfaces with a total area of 184,506 Å^2^ Top: all-atom visualization. Middle: visualization of inter-chain contacts constructed by Voronota-LT. Bottom: frequency of the chain-chain interface areas. 3D visualizations were generated using Voronota-GL.

**Table S1.** Summary of Pearson correlation coefficients between differently produced sets of areas for every input structure. We summarized every distribution of coefficients by calculating the minimal and the average values. For a more convenient display, the values were rounded up to four significant digits. Note that vanilla Voronota always uses triangulated approximations of surfaces for computing areas, while Voronota-LT uses triangulated approximations only in the additively-weighted (AW) tessellation mode.

| Regime | Method 1 | Method 2 | Pearson coeff. |  |
| --- | --- | --- | --- | --- |
|  |  |  | minimal | average |
| atom-atom area | Voronota-Vanilla-AW | Voronota-LT-Radical | 0.9609 | 0.9843 |
| atom-atom area | Voronota-Vanilla-AW | Voronota-LT-AW | 0.9990 | 1 |
| atom-atom area using uniform radii | Voronota-Vanilla-AW | Voronota-LT-Radical | 1 | 1 |
| residue-residue area | Voronota-Vanilla-AW | Voronota-LT-Radical | 0.9947 | 0.9990 |
| SAS area | Voronota-Vanilla-AW | Voronota-LT-Radical | 0.9997 | 1 |

## References

Andersen, C. W., Armiento, R., Blokhin, E., Conduit, G. J., Dwaraknath, S., Evans, M. L., Fekete, A., Gopakumar, A., Gražulis, S., Merkys, A., Mohamed, F., Oses, C., Pizzi, G., Rignanese, G.-M., Scheidgen, M., Talirz, L., Toher, C., Winston, D., Aversa, R., Choudhary, K., Colinet, P., Curtarolo, S., Di Stefano, D., Draxl, C., Er, S., Esters, M., Fornari, M., Giantomassi, M., Govoni, M., Hautier, G., Hegde, V., Horton, M. K., Huck, P., Huhs, G., Hummelshøj, J., Kariryaa, A., Kozinsky, B., Kumbhar, S., Liu, M., Marzari, N., Morris, A. J., Mostofi, A. A., Persson, K. A., Petretto, G., Purcell, T., Ricci, F., Rose, F., Scheffler, M., Speckhard, D., Uhrin, M., Vaitkus, A., Villars, P., Waroquiers, D., Wolverton, C., Wu, M., and Yang, X. (2021). OPTIMADE, an API for exchanging materials data. Sci Data, 8(1):217. [PubMed: 34385453] [PubMed Central: PMC8361091] [doi: 10.1038/s41597-021-00974-z].

Aurenhammer, F. (1987). Power Diagrams: Properties, Algorithms and Applications. SIAM J. Comput., 16(1):78–96. [doi: 10.1137/0216006].

Ben-Naim, A. (1997). Statistical potentials extracted from protein structures: Are these meaningful potentials? The Journal of Chemical Physics, 107(9):3698–3706. [doi: 10.1063/1.474725].

Bhattacharjee, S., Feng, X., Maji, S., Dadhwal, P., Zhang, Z., Brown, Z. P., and Frank, J. (2023). Time resolution in cryo-EM using a novel PDMS-based microfluidic chip assembly and its application to the study of HflX-mediated ribosome recycling. [doi: 10.1101/2023.01.25.525430].

Cazals, F. (2006). Revisiting the Voronoi description of protein-protein interfaces. Protein Sci., 15(9):2082–2092. [doi: 10.1110/ps.062245906].

Cazals, F. and Dreyfus, T. (2017). The structural bioinformatics library: modeling in biomolecular science and beyond. Bioinformatics, 33(7):997–1004. [PubMed: 28062450] [doi: 10.1093/bioinformatics/btw752].

Cazals, F., Kanhere, H., and Loriot, S. (2011). Computing the volume of a union of balls: A certified algorithm. ACM Trans. Math. Softw., 38(1):1–20. [doi: 10.1145/2049662.2049665].

Dagum, L. and Menon, R. (1998). OpenMP: an industry standard API for shared-memory programming. IEEE Comput. Sci. Eng., 5(1):46–55. [doi: 10.1109/99.660313].

Edelsbrunner, H. and Koehl, P. (2003). The weighted-volume derivative of a space-filling diagram. Proc. Natl. Acad. Sci. U.S.A., 100(5):2203–2208. [doi: 10.1073/pnas.0537830100].

Esque, J., Oguey, C., and de Brevern, A. G. (2011). Comparative analysis of threshold and tessellation methods for determining protein contacts. J Chem Inf Model, 51(2):493–507. [PubMed: 21226523] [doi: 10.1021/ci100195t].

Frenkel, D. and Smit, B. (2002). Understanding molecular simulation: from algorithms to applications. Number 1 in Computational science series. Academic Press, San Diego, 2nd ed edition.

Goede, A., Preissner, R., and Frömmel, C. (1997). Voronoi cell: new method for allocation of space among atoms: elimination of avoidable errors in calculation of atomic volume and density. J. Comput. Chem., 18:1113–1123.

Gražulis, S., Chateigner, D., Downs, R. T., Yokochi, A. F. T., Quiros, M., Lutterotti, L., Manakova, E., Butkus, J., Moeck, P., and Le Bail, A. (2009). Crystallography Open Database - an open-access collection of crystal structures. J Appl Crystallogr, 42(Pt 4):726–729. [PubMed: 22477773] [PubMed Central: PMC3253730] [doi: 10.1107/S0021889809016690].

Harder, O. F., Barrass, S. V., Drabbels, M., and Lorenz, U. J. (2023). Fast viral dynamics revealed by microsecond time-resolved cryo-EM. Nat Commun, 14(1):5649. [PubMed: 37704664] [PubMed Central: PMC10499870] [doi: 10.1038/s41467-023-41444-x].

Igashov, I., Olechnovič, K., Kadukova, M., Venclovas, C., and Grudinin, S. (2021a). VoroCNN: Deep convolutional neural network built on 3D Voronoi tessellation of protein structures. Bioinformatics, page btab118. [PubMed: 33620450] [doi: 10.1093/bioinformatics/btab118].

Igashov, I., Pavlichenko, N., and Grudinin, S. (2021b). Spherical convolutions on molecular graphs for protein model quality assessment. Mach. Learn.: Sci. Technol., 2(4):045005. [doi: 10.1088/2632-2153/abf856].

Imai, H., Iri, M., and Murota, K. (1985). Voronoi Diagram in the Laguerre Geometry and Its Applications. SIAM J. Comput., 14(1):93–105. [doi: 10.1137/0214006].

Kim, D. S., Kim, D., and Cho, Y. (2005). Euclidean Voronoi diagrams of 3D spheres: Their construction and related problems from biochemistry. Lect. Notes. Comput. Sc., 3604:255–271.

Klenin, K. V., Tristram, F., Strunk, T., and Wenzel, W. (2011). Derivatives of molecular surface area and volume: simple and exact analytical formulas. J. Comput. Chem., 32(12):2647–2653. [PubMed: 21656788] [doi: 10.1002/jcc.21844].

Lee, M., Sugihara, K., and Kim, D.-S. (2022). Robust Construction of Voronoi Diagrams of Spherical Balls in Three-Dimensional Space. Comput. Aided Design, 152:103374. [doi: 10.1016/j.cad.2022.103374].

Lensink, M. F., CAPRI-participants, and Wodak, S. J. (2023). Impact of AlphaFold on structure prediction of protein complexes: The CASP15-CAPRI experiment. Proteins, 91(12):1658–1683. [PubMed: 37905971] [doi: 10.1002/prot.26609].

Li, A. J. and Nussinov, R. (1998). A set of van der Waals and coulombic radii of protein atoms for molecular and solvent-accessible surface calculation, packing evaluation, and docking. Proteins, 32(1):111–127. [PubMed: 9672047].

Lu, J., Lazar, E. A., and Rycroft, C. H. (2023). An extension to Voro++ for multithreaded computation of Voronoi cells. Computer Physics Communications, 291:108832. [doi: 10.1016/j.cpc.2023.108832].

McConkey, B. J., Sobolev, V., and Edelman, M. (2002). Quantification of protein surfaces, volumes and atom-atom contacts using a constrained Voronoi procedure. Bioinformatics, 18(10):1365–1373. [PubMed: 12376381].

McConkey, B. J., Sobolev, V., and Edelman, M. (2003). Discrimination of native protein structures using atom-atom contact scoring. Proc. Natl. Acad. Sci. U.S.A., 100(6):3215–3220. [PubMed: 12631702] [PubMed Central: PMC152272] [doi: 10.1073/pnas.0535768100].

Neira, J. L., Davis, B., Ladurner, A. G., Buckle, A. M., Gay, G. d. P., and Fersht, A. R. (1996). Towards the complete structural characterization of a protein folding pathway: the structures of the denatured, transition and native states for the association/folding of two complementary fragments of cleaved chymotrypsin inhibitor 2. Direct evidence for a nucleation-condensation mechanism. Fold Des, 1(3):189–208. [PubMed: 9079381] [doi: 10.1016/s1359-0278(96)00031-4].

Olechnovič, K., Valančauskas, L., Dapkūnas, J., and Venclovas, C. (2023). Prediction of protein assemblies by structure sampling followed by interface-focused scoring. Proteins, 91(12):1724–1733. [PubMed: 37578163] [doi: 10.1002/prot.26569].

Olechnovič, K. and Venclovas, C. (2014). Voronota: A fast and reliable tool for computing the vertices of the Voronoi diagram of atomic balls. J. Comput. Chem., 35(8):672–681. [PubMed: 24523197] [doi: 10.1002/jcc.23538].

Olechnovič, K. and Venclovas, C. (2017). VoroMQA: Assessment of protein structure quality using interatomic contact areas. Proteins, 85(6):1131–1145. [PubMed: 28263393] [doi: 10.1002/prot.25278].

Olechnovič, K. and Venclovas, C. (2021). VoroContacts: a tool for the analysis of interatomic contacts in macromolecular structures. Bioinformatics, page btab448. [PubMed: 34132767] [doi: 10.1093/bioinformatics/btab448].

Olechnovič, K. and Venclovas, C. (2023). VoroIF-GNN: Voronoi tessellation-derived protein-protein interface assessment using a graph neural network. Proteins, 91(12):1879–1888. [PubMed: 37482904] [doi: 10.1002/prot.26554].

Poupon, A. (2004). Voronoi and Voronoi-related tessellations in studies of protein structure and interaction. Curr. Opin. Struct. Biol., 14(2):233–241. [PubMed: 15093839] [doi: 10.1016/j.sbi.2004.03.010].

Rose, Y., Duarte, J. M., Lowe, R., Segura, J., Bi, C., Bhikadiya, C., Chen, L., Rose, A. S., Bittrich, S., Burley, S. K., and Westbrook, J. D. (2021). RCSB Protein Data Bank: Architectural Advances Towards Integrated Searching and Efficient Access to Macromolecular Structure Data from the PDB Archive. J Mol Biol, 433(11):166704. [PubMed: 33186584] [PubMed Central: PMC9093041] [doi: 10.1016/j.jmb.2020.11.003].

Schweke, H., ELIXIR-participants, and Wodak, S. J. (2023). Discriminating physiological from non-physiological interfaces in structures of protein complexes: A community-wide study. Proteomics, 23(17):e2200323. [PubMed: 37365936] [doi: 10.1002/pmic.202200323].

Stenqvist, B., Thuresson, A., Kurut, A., Vacha, R., and Lund, M. (2013). Faunus -a flexible framework for Monte Carlo simulation. Molecular Simulation, 39(14-15):1233–1239. [doi: 10.1080/08927022.2013.828207].

Todhunter, I. (1863). Spherical trigonometry, for the use of colleges and schools: with numerous examples. Macmillan.

UniProt Consortium (2023). UniProt: the Universal Protein Knowledgebase in 2023. Nucleic Acids Res, 51(D1):D523–D531. [PubMed: 36408920] [PubMed Central: PMC9825514] [doi: 10.1093/nar/gkac1052].

Varadi, M., Anyango, S., Deshpande, M., Nair, S., Natassia, C., Yordanova, G., Yuan, D., Stroe, O., Wood, G., Laydon, A., Zidek, A., Green, T., Tunyasuvunakool, K., Petersen, S., Jumper, J., Clancy, E., Green, R., Vora, A., Lutfi, M., Figurnov, M., Cowie, A., Hobbs, N., Kohli, P., Kleywegt, G., Birney, E., Hassabis, D., and Velankar, S. (2022). AlphaFold Protein Structure Database: massively expanding the structural coverage of protein-sequence space with high-accuracy models. Nucleic Acids Res, 50(D1):D439–D444. [PubMed: 34791371] [PubMed Central: PMC8728224] [doi: 10.1093/nar/gkab1061].

Verdonk, M. L., Ludlow, R. F., Giangreco, I., and Rathi, P. C. (2016). Protein-Ligand Informatics Force Field (PLIff): Toward a Fully Knowledge Driven “Force Field” for Biomolecular Interactions. J. Med. Chem., 59(14):6891–6902. [PubMed: 27353137] [doi: 10.1021/acs.jmedchem.6b00716].

Voronoi, G. (1908). Nouvelles applications des parametres continus a la theorie des formes quadratiques. J. Reine Angew. Math., 134:198–287.

wwPDB consortium (2019). Protein Data Bank: the single global archive for 3D macromolecular structure data. Nucleic Acids Res, 47(D1):D520–D528. [PubMed: 30357364] [PubMed Central: PMC6324056] [doi: 10.1093/nar/gky949].

Xie, T. and Grossman, J. C. (2018). Crystal Graph Convolutional Neural Networks for an Accurate and Interpretable Prediction of Material Properties. Phys. Rev. Lett., 120(14):145301. [doi: 10.1103/PhysRevLett.120.145301].

